# Age-related degradation of behavioral and network features of *Aplysia* escape locomotion

**DOI:** 10.1101/2025.09.30.679365

**Authors:** Viral K. Mistry, Daniel Martinez, William Frost

## Abstract

*Aplysia californica* has been a useful model system for studies of the neural basis of behavior, learning, and aging. While the latter topic has been explored with respect to several of its simple reflex behaviors, this study represents the first examination of how one of Aplysia’s more complex behaviors, escape locomotion, is affected in animals nearing the end of their natural lifespan. Old animals (12-13mo) showed a greatly reduced gallop response compared with middle-aged adults (5-7mo), together with a loss of locomotion onset latency sensitization. Large-scale VSD imaging was used to record motor programs in isolated brain preparations from middle-aged vs. elderly animals. Old brains displayed the same loss of onset latency sensitization seen in the intact old animal behavior, and also a reduced number of cycles per locomotion episode. Brains from middle-aged animals showed an unchanged number of motor program cycles from that observed in intact animals, but a much more transient motor program onset latency sensitization. A further age-related finding was that while in middle-aged brains repeatedly eliciting the motor program led to progressively increasing cumulative activity across trials, in old brains this same procedure led to progressively decreasing activity. Some of our results are consistent with peripheral processes working in concert with the CNS as animals age to support healthy locomotion behavior and its modification by learning, or with early changes in the brain that are not yet expressed in behavior.

## INTRODUCTION

Aging results in progressive decline in normal physiological function across species [1, 2]. In the nervous system, aging causes cellular and molecular changes that result in impaired behavioral function and reduced memory formation. This includes reduced synaptic plasticity, reduced neurotransmitter expression, the loss of neurons, and reduced circuit activity [3]. The longevity and complexity of vertebrates creates an opportunity for the use of simpler invertebrates for studying the neural correlates of age-related behavioral and cognitive decline, with the goal of revealing evolutionarily conserved principles of aging [4, 5]. Investigations into *C. elegans* and *Drosophila melanogaster*, for instance, have taken advantage of these species’ simpler physiology, nervous systems, and behaviors to provide new insights into the cellular and molecular mechanisms of aging [6–11]. However, there remains a need for simple models that can address the larger network-level components of behavioral and cognitive aging and test potential interventions for rescuing decline.

*Aplysia californica* is an invertebrate gastropod mollusk with a repertoire of behaviors that have been the basis of many foundational investigations into the neural basis of behavior and learning [12, 13]. *Aplysia* also undergoes progressive behavioral and neurophysiological decline as it ages [14, 15], and can be acquired from a facility where they are raised from eggs to adulthood under controlled conditions [16]. Aged *Aplysia* display cellular, molecular, and behavioral changes that have made them a valuable model system for studying age-related behavioral and cognitive decline [17–24]. However, these previous studies have largely focused on *Aplysia’s* simple reflexes, such as its tail withdrawal or gill-siphon withdrawal reflex, which involve only a small network of identified neurons that produce behaviors that only persist for a few seconds. *Aplysia* also possesses a more complex behavior, escape locomotion, which has two phases - an initial fast “gallop” phase which transitions into a slower “crawl” phase [25]. The behavior arises from a low-dimensional spiral attractor network state and is comprised of neuronal ensembles that are spatially and functionally clustered within the pedal ganglia [26, 27]. A central pattern generator generates this behavior by driving a large population of efferent neurons to rhythmically burst for many minutes, producing a repeating wave of head-to-tail muscle contraction that propels the animal forward [28, 29]. The motor network is known to be primarily driven by serotonin with key modulatory neurons located in the pedal and cerebral ganglia [30–32]. Exploration into learning in this behavior, however, has been limited to a few studies [33, 34], and no prior study has explored how said learning in this behavior is affected by aging. This presents an opportunity to investigate how aging affects both naive motor function and memory formation in a more complex behavior within a simple tractable nervous system. This behavior, and the network that produces it, has potential to generate insights into behavioral and cognitive aging at a network level that may have potential relevance to investigations across species.

In this study, we demonstrate for the first time that elderly *Aplysia* display behavioral and cognitive aging of specific parameters of their escape locomotion behavior and network. Aged *Aplysia* lose the ability to behaviorally sensitize locomotion onset and show significant decline in their galloping capacity. Within their isolated brains, the deficits are even more significant, demonstrating that these effects of aging on escape behavior are driven by CNS aging, and suggesting the presence of compensatory mechanisms for CNS aging and early cognitive decline in the brain that does not yet manifest in the behavior. Elderly *Aplysia* brains also responded less vigorously to an initial stimulus than middle-aged adult brains. With repeated stimulation, we observed inverse effects with respect to age – successive stimuli produced increased overall neuronal activity during the motor program in middle-aged brains, but produced decreased overall neuronal activity in old brains. We also show that the resting state activity of elderly *Aplysia*’s motor network is reduced with age, suggesting broader network degeneration that may explain these behavioral and cognitive deficits.

These findings demonstrate that aging has distinct and heterogeneous effects on the *Aplysia* escape locomotion network and validate it as a model for further investigations into the cellular, molecular, and network dynamics of behavioral and cognitive aging. This study will also inform future work to investigate methods of mitigating or rescuing this decline.

## METHODS

*Preparation – Aplysia californica* of desired ages were obtained from the National Resource for *Aplysia* at the University of Miami’s Rosenstiel School of Marine, Atmospheric, and Earth Science (Coral Gables, FL). Animal hatch date was provided with each shipment. Middle-aged adult *Aplysia* were 5-7mo, and weighed between 21-40g. Elderly adult *Aplysia* were 12-13mo and weighed between 100-250g. Animals were maintained in chilled (13.5-16.5°C) recirculating artificial seawater (ASW) systems prior to experiments. The ASW was made using Instant Ocean mix. Experiments involving isolated brains were performed in filtered ASW.

*Behavioral experiments - Aplysia* were weighed and then placed alone into the top tier of a plexiglass recirculating artificial seawater tank. 1 hour after being moved into the testing region, locomotion was elicited by applying 1mL of 5M NaCl to the tail of the animal. Locomotion was filmed briefly before salt application and then for 2 minutes following the beginning of salt application via a digital camcorder that was perpendicular to the tank, allowing for full view of the lateral profile of the entire animal. Videos were analyzed and hand-scored for behavioral features using ShotCut. Head reach latency was measured as the time between the beginning of the application of the NaCl solution to the maxima of the first head extension. Gallops and crawls were distinguished as previously defined in [25, 35]: gallop cycles required an arched head reach followed by a simultaneous mid-body & tail pull, while crawl cycles see distinct head reach followed by a mid-body slide and then a tail pull. As the animals side profiles were used, distance traveled could not be measured as animals were not consistently locomoting in a straight line.

*Optical recordings -* In experiments where we sought to characterize the efferent locomotion network, we optically recorded voltage activity using a fast voltage-sensitive dye to record the action potentials of rhythmically active gallop/crawl neurons in the pedal ganglion, as in [26, 27, 35]. *Aplysia’s* central ganglia, consisting of the cerebral ganglion connected to the left & right pleural and pedal ganglia were dissected and pinned to the bottom of a Sylgard (Dow Corning) lined Petri dish containing filtered ASW. During the entire experiment, the temperature was maintained between and 14.0°C and 15.0 °C by passing filtered ASW through a feedback-controlled in-line Peltier cooling system (Model SC-20, Warner Instruments) using a peristaltic pump (Model 720, Instech Laboratories). Temperature was monitored with a BAT-12 thermometer fitted with an IT-18 microprobe (Physitemp Instruments). Pedal ganglion neurons were stained by periodically applying pressure to a PE tube filled with 0.2mg/mL RH-155 (Toronto Research Chemicals, Toronto, CA), pressed against the surface of the ganglion for 1 hour. Following staining, the pedal ganglion was transferred and pinned to a Slygard-lined perfusion chamber used for optical recording, where the PdN9, a tail nerve, contralateral to the stained pedal ganglion was pulled into a suction electrode. To improve visualization, two small pieces of silicone were placed in the chamber on opposite sides of the stained ganglion, allowing for a glass coverslip fragment to be pressed down and held in place to flatten the surface of the stained pedal ganglion (as described in [35]). The preparation was trans-illuminated by a 735nm LED (Thor Labs), and the light was collected by a 10x 0.6 NA water-immersion objective lens (Olympus) and passed through a phototube to reach either the photodiode array or a parfocal camera (Optronics). The preparation rested on the optical imaging rig for 60 minutes following nerve suction before acquisitions began. Fictive escape locomotion motor programs were elicited via stimulation of the contralateral PdN9 (8V, 20Hz, 2.5s stimulus train, 5ms pulses). Voltage activity was captured using a RedShirtImaging PDA-III photodiode array consisting of 464 photodiodes sampled at 1600 Hz. Data were acquired using RedShirtImaging’s Neuroplex software. Optical data were AC-coupled to zero the baseline of the traces and then amplified 100x by the PDA-III. After acquisition, optical data were band-pass filtered in the Neuroplex software (Butterworth, 5Hz high pass, 100Hz low pass) and saved as text files. Independent component analysis was run on the optical data in MATLAB (Mathworks), as made available in [36], to obtain action potentials from individual neurons. Following ICA, the components had a manual threshold applied to convert the action potential spike trains into binary spike times for further analysis of neuronal activity.

*Analysis of spike trains generated via optical recordings* – Spike times were calculated from the binary spike train files generated from the optical recordings. Neural correlates of behavior (onset latency and cycle number) were calculated using a custom MATLAB script to determine the time from stimulus onset to the termination of the first rhythmic burst (onset latency) as well as the number of rhythmic bursts within the first two minutes from stimulus onset (cycle number). Spike times were used to generate spike frequency in 1-second time bins for each neuronal trace and averaged for each recording to generate average spike frequency. An unsupervised consensus clustering algorithm described in [26] was used to determine functional ensembles in the motor program and to identify candidate rhythmic ensembles in recordings of spontaneous activity. Custom MATLAB script was used to determine rhythmicity within an averaged ensemble: Welch’s method was used to calculate power spectral density and spectral peaks with adaptive prominence were identified. Peak power ratio (max peak power divided by mean power), coefficient of variation of the spectrum, and spectral entropy were used in the following equation to generate a “rhythmicity score”:

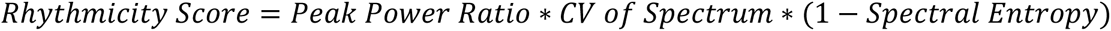

A positive rhythmicity score indicated a rhythmic cluster. If a cluster was determined to be rhythmic, the burst duration, intraburst frequency, and cycle frequency for every burst performed by the cluster during the recording were then calculated. The mean burst duration, intraburst frequency, and cycle frequency were then calculated for all the rhythmic clusters within a single recording.

## RESULTS

### Elderly Aplysia cannot sensitize onset latency and lose galloping capacity

To determine the effects of aging on *Aplysia’s* escape locomotion behavior and capacity for learning, we developed a same-site behavioral non-associative learning protocol for this behavior that we then applied to both middle-aged adult (5-7mo) and elderly adult *Aplysia* (12-13mo). In this behavioral paradigm, we provided a repeated same-site aversive salt stimulus (1mL 5M NaCl) to the animal’s tail. In the experimental group (n = 16 for both middle-aged and old animals), we provided 5 stimuli 10 min apart. In the control group (n = 16 for both middle-aged and old animals), we provided 2 stimuli 40 min apart – the equivalent amount of time as the experimental group (**Fig. 1A**). We then analyzed various parameters of the behavior to identify potential behavioral features of either sensitization (increased response) or habituation (decreased response). We chose three behavioral parameters (head reach latency, cycle number within 2 minutes of the stimulus, and the percentage of locomotion cycles that are gallops) that were easily quantifiable and represented different features of locomotion to see if age-related decline differentially affected the overall behavior. Head reach latency represented locomotion onset, cycle # represented duration, and the share of cycles that were gallop vs. crawl represented intensity.

**Figure 1.**
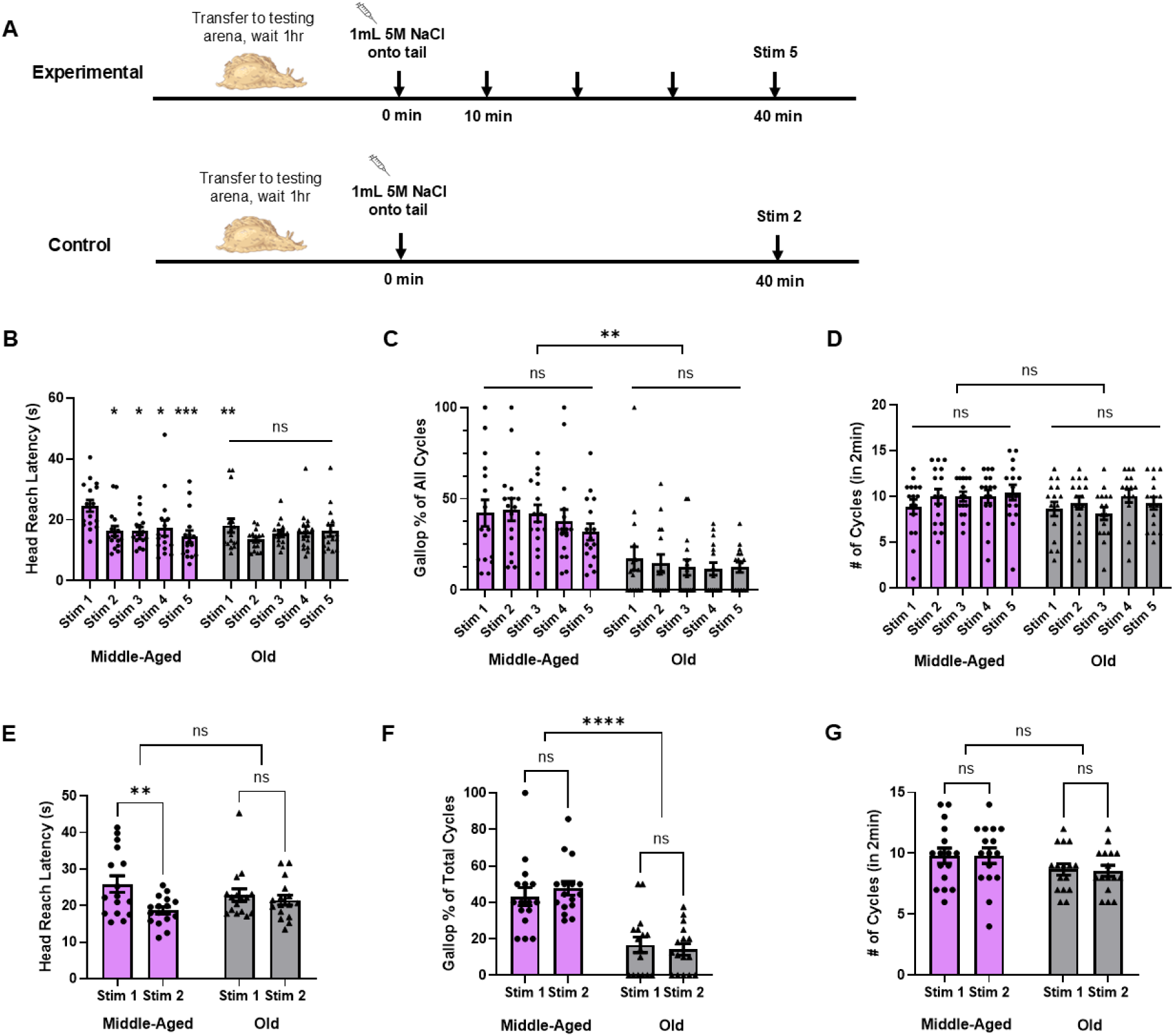
Aging causes decline in specific parameters of escape locomotion behavior. **A)** Behavioral protocol used in middle-aged and old *Aplysia*, with experimental group (n = 16 for middle-aged and old) receiving 5 stimuli at 10 min ISI and control group (n = 16 for middle-aged and old) receiving 2 stimuli at 40 min ISI (equivalent time as experimental). **B)** Head reach latency significantly quickened after an initial stimulus in middle-aged *Aplysia* and remained quickened with successive stimui (* for *p* < 0.05, *** for *p* < 0.001). Head reach latency to stim 1 in middle-aged *Aplysia* is also significantly slower than latency to stim 1 in old *Aplysia*. **C)** Gallop share of all locomotion cycles was unchanged within middle-aged and old groups, but was significantly elevated in middle-aged versus old (** for *p* < 0.01). **D)** No change in cycle number produced within 2 minutes in either middle-aged or old *Aplysia*. **E)** Head reach latency is significantly quickened in middle-aged *Aplysia* in control condition, demonstrating persistent sensitization. **F)** Gallop share in control condition demonstrates again only age effect, not effect from stimulation. **G)** Cycle number in control condition shows no effect from stimulation or between groups.

In the experimental group, a RM two-way ANOVA showed a significant effect of repeated stimulation on head reach latency (F(4, 75) = 3.859, *p* = 0.007) (**Fig. 1B)**. Multiple comparisons with Tukey’s post-hoc tests revealed a quickening of head reach latency from stim 1 to stim 2 in middle-aged *Aplysia* (stim 1: 24.6 ± 1.9s vs. stim 2: 16.3 ± 1.6s, *p* = 0.01). This quickening of head reach latency persisted with each follow-up stimulus for the middle-aged age group (stim 3: 16.3 ± 1.3s, *p* = 0.01; stim 4: 16.6 ± 2.6s, *p* = 0.03; stim 5: 14.6 ± 1.9s, *p* = 0.0009). By comparison, there was no change in head reach latency from stim 1 to stim 2 in old *Aplysia* (stim 1: 18.1 ± 2.3s vs. stim 2: 13.5 ± 0.8s, *p* = 0.37), or in any follow up stimuli. Head reach latency to the first stimulus was also significantly different between middle-aged and old *Aplysia* in the experimental group (*p* = 0.008). In the control group, a RM two-way ANOVA also revealed a significant effect from the stimuli (F(1, 15) = 9.911, *p* = 0.007) (**Fig. 1E)**. Tukey’s post-hoc tests also revealed a significant quickening of head reach latency in the control group from stim 1 to stim 2 in middle-aged *Aplysia* (stim 1: 25.9 ± 2.3s vs. stim 2: 18.7 ± 0.9s, *p* = 0.004), but no change in head reach latency from stim 1 to stim 2 in old *Aplysia* (stim 1: 22.9 ± 1.7s vs. stim 2: 21.5 ± 1.3s, *p* = 0.85). These results reveal an identified cognitive feature of *Aplysia*’s locomotion behavior – sensitization of head reach latency – that is lost with aging. This behavioral sensitization in middle-aged animals persisted with repeated stimuli and as shown by the control group, lasted after a single trial for at least 40 minutes.

The experimental group observed no change in gallop share over repeated stimulation within either the middle-aged or old groups (F(4, 75) = 0.77, *p* = 0.55) (**Fig. 1C)**. There was, however, a significant difference between the middle-aged and old groups in gallop share (F(1, 75) = 57.91, *p* < 0.0001, mean of middle-aged: 39.6% vs. mean of old: 13.8%). The control group displayed the same phenomenon: no effect from stimulation within either age group (F(1, 15) = 0.1055, *p* = 0.75), but a difference between the age groups (F(1, 15) = 37.04, *p* <0.0001, mean of middle-aged: 45.4% vs. mean of old: 15.5%) (**Fig. 1F)**. Therefore, this stimulus protocol did not produce learning-induced changes in galloping in middle-aged or old animals and instead revealed an age-based behavioral loss in galloping that could not be rescued by repeated stimulation.

Meanwhile the number of cycles performed within the first 2 minutes following the stimulus in the experimental group was unchanged by repeated stimulation (F(4, 75) = 1.021, *p* = 0.40) or by age (F(1,75) = 3.160, *p* = 0.08, mean of middle-aged: 9.9 cycles vs. mean of old: 9.1 cycles) (**Fig. 1D)**. In the control group we also found cycle number did not change from follow-up stimulation (F(1, 15) = 0.0134, *p* = 0.91) or between age groups (F(1,15) = 3.871, *p* = 0.07, mean of middle-aged: 9.8 cycles vs. mean of old: 8.6 cycles) (**Fig. 1G)**. Thus, this behavioral parameter did not develop any learning induced changes via this behavioral protocol in either age group, and did not display any significant age-related decline.

Here, we demonstrated that aging is associated with a loss of sensitization learning in a specific parameter of locomotion – head reach latency. A different parameter, gallop share, was unchanged by learning but was diminished with aging. Yet another parameter, cycle number, was unchanged by both learning and aging. Aging thus degrades both motor gallop share and cognitive abilities in *Aplysia’s* locomotion behavior.

### Isolated brains from old animals also cannot sensitize onset latency and show reduced cycling in the fictive motor program

After we identified various parameters of behavioral locomotion that differentially undergo learning or behavioral loss with age, we then sought to identify neural correlates of these behavioral deficits. With this, we sought to determine if these changes were the result of CNS changes alone or whether they required the periphery. In *Aplysia*, the locomotion efferent neurons are located in the pedal ganglia [29], and previous work has shown that a fictive motor program can be generated in an isolated brain via a stimulus to the tail nerve pedal nerve 9 (PdN9) [26, 32, 35]. To identify how aging specifically affects the *Aplysia* brain, we induced fictive locomotion in isolated *Aplysia* brains (middle-aged and old) via a stimulus to PdN9. As with the intact animals, in the experimental group (n = 6 for both middle-aged and old brains), we gave 5 stimuli 10 minutes apart, and in the control group (n = 6 for both middle-aged and old brains), we gave 2 stimuli 40 minutes apart (**Fig. 2A)**. We recorded the ensuing fictive motor program using voltage-sensitive dye imaging and extracted the true spiking activity of dozens of pedal efferent neurons per preparation to characterize the neural correlates of two locomotor parameters – onset latency (equivalent of head reach latency) and the number of locomotion cycles generated in 2 minutes.

**Figure 2.**
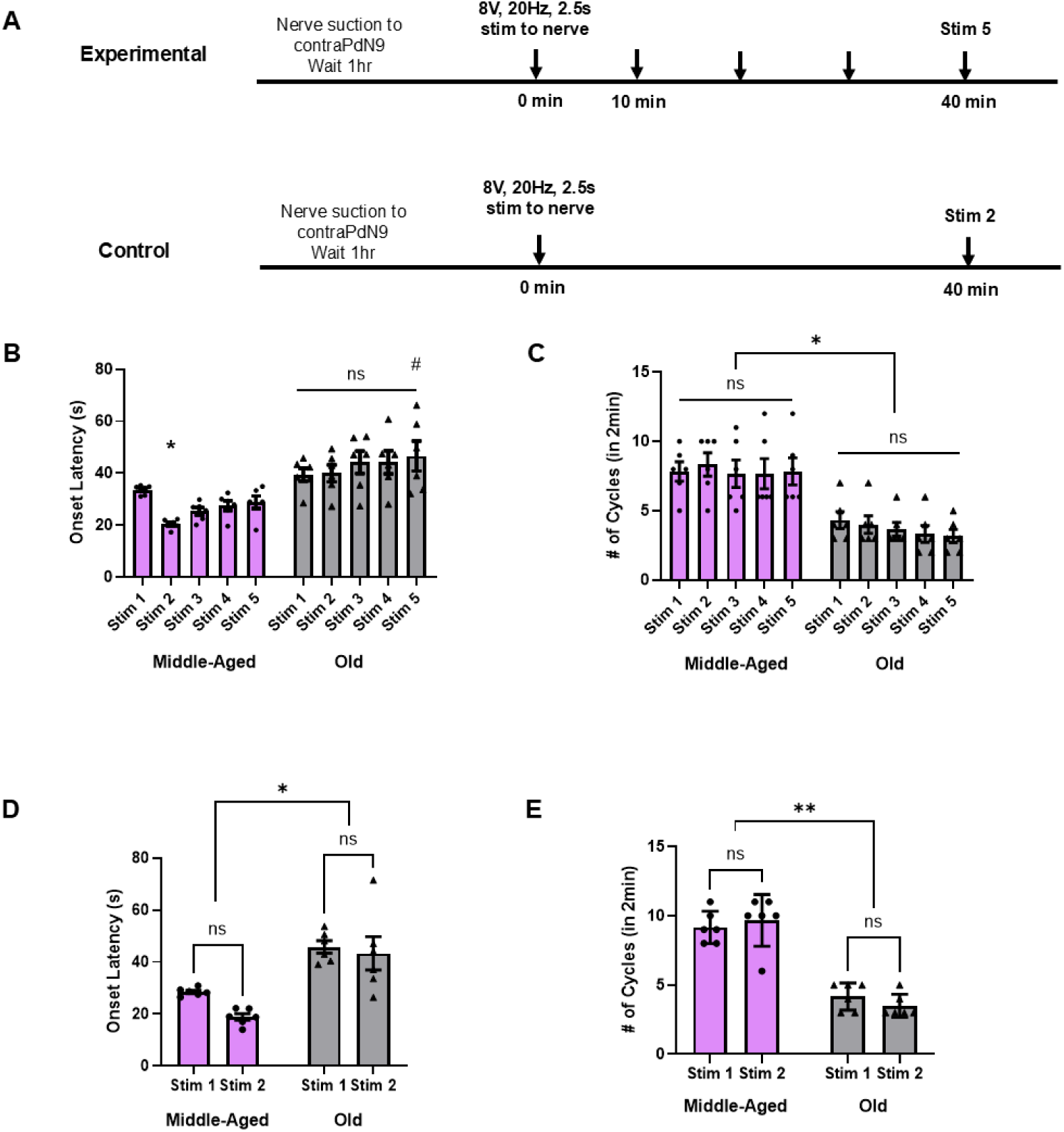
Aging also produces deficits in fictive locomotion in isolated brains. **A)** Protocol for stimulation in isolated brains, with the experimental group (n = 6 for middle-aged and old) receiving 5 stimuli to PdN9 at 10 min ISI, and control group (n = 6 for middle-aged and old) 2 stimuli at 40 min ISI (equivalent time as experimental). **B)** Onset latency only shows quickening from stim 1 to stim 2 in middle-aged brains (* for *p* < 0.05). Old brains show no change in onset latency with repeated stimulation. Onset latency to stim 5 in old brains is significantly slower than stim 5 in middle-aged brains (# for *p* < 0.05). **C)** Cycle number is unchanged in both middle-aged and old brains with repeated stimulation, but is significantly reduced at every stimulus in old brains compared to middle-aged brains (* for *p* < 0.05). **D)** Onset latency in control experiments shows no quickening of onset latency in either middle-aged or old brains at follow up stimulus, but old brains are significantly slower at both stimuli (* for *p* < 0.05). **E)** Cycle number in control experiments demonstrates no change with stimulus but significant decline with age (* for *p* < 0.05).

In the experimental group, a RM two-way ANOVA identified significant effects on onset latency from repeated stimuli (F(4, 20) = 5.346, *p* = 0.004), and age (F(1, 5) = 18.68, *p* = 0.008) (**Fig. 2B)**. Multiple comparisons with Tukey’s post-hoc tests revealed a significant quickening in onset latency from stim 1 to stim 2 in middle-aged isolated brains (stim 1: 33.6 ± 0.7s vs. stim 2: 20.4 ± 0.7s, *p* = 0.03). However, differing from our behavioral results, this quickening faded by stim 3 (25.4 ± 1.4s, *p* = 0.42), and remained unchanged by stim 5 (28.8 ± 2.4s, *p* = 0.93), suggesting latency sensitization is more transient in a middle-aged isolated brain compared to the intact animal. In old isolated brains, by comparison, there was no sensitization of onset latency from stim 1 to stim 2 (stim 1: 39.4 ± 2.5s vs. stim 2: 39.9 ± 3.2s, *p* > 0.99), nor any significant change by stim 5 (46.6 ± 5.7s, *p* = 0.58), once again demonstrating a loss of sensitization learning with aging. In the control group, a RM two-way ANOVA only identified a significant difference in onset latency from age (F(1, 5) = 35.73, *p* = 0.002), with no sensitization effect persisting across the 18 min stimulus gap (F(1,5) = 4.636, *p* = 0.08) (**Fig. 2D)**. Multiple comparisons with Tukey’s tests in the control data showed no change in onset latency from stim 1 to stim 2 in middle-aged (stim 1: 28.5 ± 0.6s vs. stim 2: 18.9 ± 1.2s*, p* = 0.17) or old isolated brains (stim 1: 45.9 ± 2.4s vs. stim 2: 43.3 ± 6.4s, *p* = 0.91). However, there was a significant difference in onset latency between middle-aged and old brains at stim 1 (*p* = 0.02) and stim 2 (*p* = 0.04). Thus, while middle-aged isolated brains demonstrated onset latency sensitization, it did not persist across the 5 trials as it did in behavior. Old isolated brains display a loss of onset latency sensitization, and respond to tail nerve stim with fewer cycles, suggesting an earlier age-related decline in the CNS than observed in the intact animal.

Meanwhile, cycle number in the experimental group was unchanged by repeated stimulation (F(4,20) = 1.000, *p* = 0.43), but was significantly reduced in the aged brains (F(1,5) = 38.89, *p* = 0.002, mean of middle-aged: 7.9 cycles vs. mean of old: 3.7 cycles) **(Fig. 2C)**. This was also seen in the control group, where there was no effect on cycle number from follow-up stimulation (F (1,5) = 0.033, *p* = 0.86) but there was a significant effect from age (F(1,5) = 61.49, *p* = 0.0005, mean of middle-aged: 9.4 cycles vs. mean of old: 3.8 cycles) (**Fig. 2E)**. Thus, while cycle number was unchanged within each age group with repeated stimulation, isolated brains from old animals generated far fewer cycles than those from middle-aged animals.

These results demonstrate that age-related behavioral and cognitive decline is also present in *Aplysia*’s CNS, and that these deficits are even more severe than what was observed in the intact animal behavior. This raises the possibility that early markers of aging are emerging in the isolated brain that are not observed in the whole animal.

### Repeated stimulation makes middle-aged networks more active and old networks less active

In addition to allowing us to characterize how age affected onset latency and cycling in the fictive motor program, the voltage-sensitive dye imaging technique described in the previous section also allowed us to assess whether age affects the activity of the pedal ganglion neurons as a population during the fictive motor program. Our behavioral experiments identified a significant decline in galloping with age, which represents a more vigorous and intensive form of locomotion than crawling. We were interested in determining if aging also caused the locomotion network to become less vigorous by reducing overall population activity after a stimulus. To determine this, we calculated the average spike frequency over time in each VSD experiment described in the previous section (n = 6 for both middle-aged and old brains), to determine if overall neuronal activity changed due to age or repeated stimulation within the first 2 minutes after a stimulus.

We compared changes in neuronal activity between our middle-aged and old groups by determining the average spike frequency for each VSD preparation and comparing changes in the average number of spikes in a neuron across the entire recording (**Fig. 3C-D)**. In our experimental group, a two-way ANOVA found significant effects from repeated stimulation on average neuronal activity over time (F(4,50) = 5.582, *p* = 0.0009) and age (F(1,50) = 298.4, *p* < 0.0001) (**Fig. 3E)**. Tukey’s tests found significant increases in the average number of spikes per neuron in the middle-aged group from stim 1 to stim 3 (stim 1: 106 ± 2.8 vs. stim 3: 120.0 ± 2.0, *p* = 0.01), stim 4 (126.7 ± 2.5, *p* < 0.0001), and stim 5 (127.1 ± 2.5, *p* < 0.0001). This indicated that the middle-aged brains were producing progressively stronger fictive locomotion with each successive stimulus, an example of sensitization. In our old experimental group, by comparison, we observed a steady decrease in activity with repeated stimulus trials. Tukey’s post-hoc tests identified significant decreases in the average # of spikes/cell in the old group from stim 1 to stim 2 (stim 1: 114.9 ± 3.5 vs stim 2: 100 ± 3.1, *p* = 0.01), stim 3 (88 ± 2.0, *p* < 0.0001), stim 4 (74.4 ± 2.1, *p* < 0.0001), and stim 5 (74.7 ± 2.1, *p* < 0.0001). Thus, old brains were producing progressively weaker fictive locomotion with successive stimuli, an example of habituation. In our control group (**Fig. 3F)**, a two-way ANOVA revealed no effect from stimulation (F(1, 20) = 0.8676, *p* = 0.36), only an effect from age (F(1, 20) = 588.5, *p* < 0.0001, mean of middle-aged: 130.5 vs. mean of old: 70.6), indicating that old brains in general produce weaker fictive locomotion, a clear effect of aging (**Fig. 3G)**. These results demonstrate that middle-aged brains produce more intensive locomotion that sensitizes with repeated stimulation. In old brains, however, the network starts off weaker and habituates with repeated stimulation.

**Figure 3.**
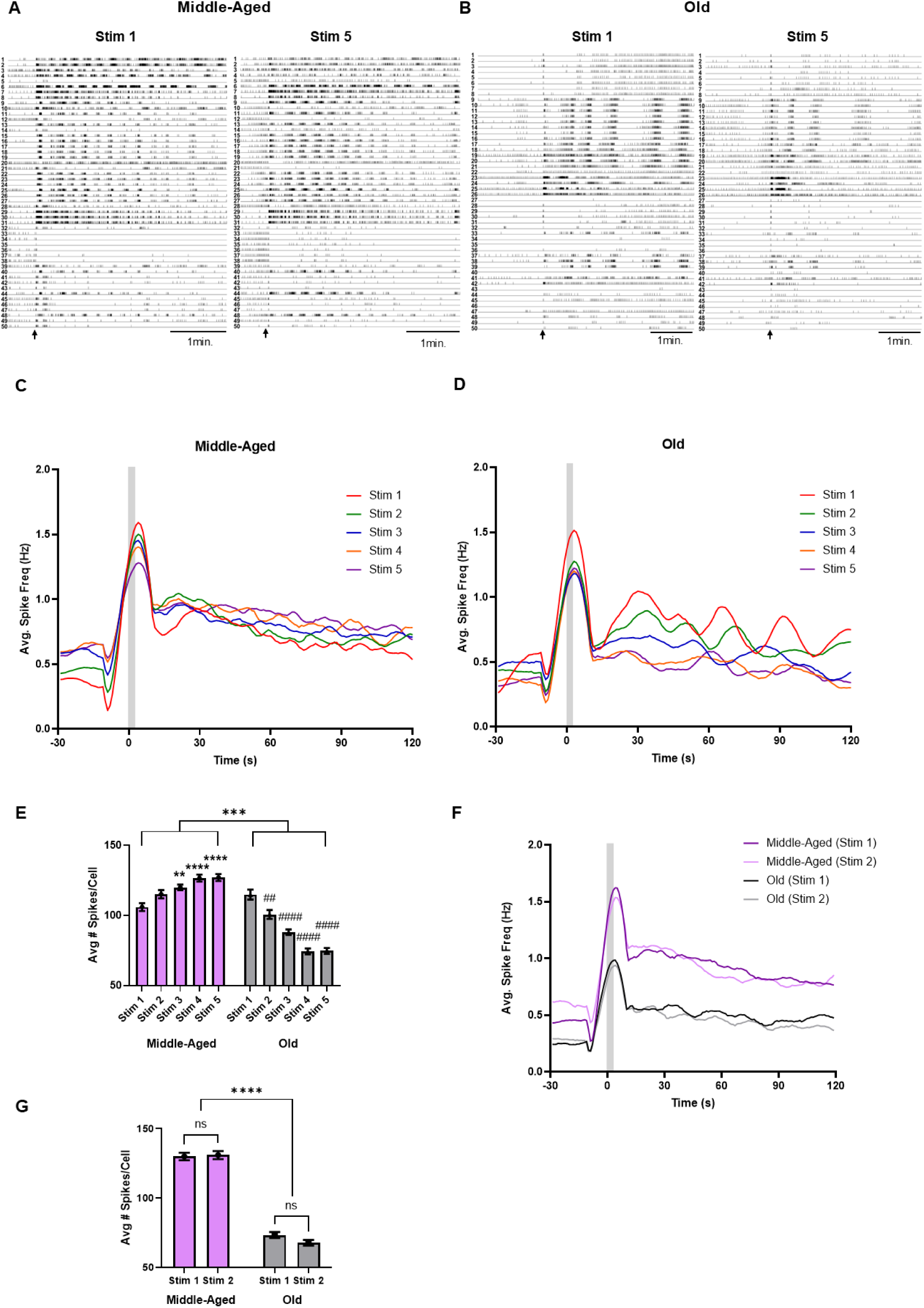
Aging reduces neural activity following a stimulus. **A)** Example of spike rasters generated from VSD imaging in the same middle-aged isolated brain at stim 1 (left) and stim 5 (right). First 50 neurons identified with ICA displayed. Each numbered neuron in stim 1 is the same neuron in stim 5. Stimulus indicated with black arrow. **B)** Same type of raster demonstrated in **A**, but from an old brain. **C)** Averaged spike frequency graphs for all recordings at each experimental stimulus in middle-aged brains. Rolling 10-s average spike frequency plotted. **D)** Same type of graph as **C**, but with old brains under experimental condition. **E)** Average spikes/cell significantly increased with followup stimuli in middle-aged isolated brains (** for *p* < 0.01, *** for p < 0.001). In comparison, Average spikes/cell significantly decreased with follow-up stimuli in old isolated brains (## for *p* < 0.01, #### for *p* < 0.0001). **F)** Average spike frequency graphs for control recordings in middle-aged (purple) and old (grey) isolated brains. Rolling 10-s average spike frequency plotted. **G)** Average spikes/cell did not change in control condition within either middle-aged or old brains, but was significantly reduced in old brains (**** for *p* < 0.000).

The network activity changes in both middle-aged and old brains, but are these changes occurring in a consistent fashion between repeated motor programs? To answer this, we tested if a single curve could fit all the trials in middle-aged or old recordings. We found that a single two-phase decay curve could fit all the spike frequency data in the middle-aged experimental group (r^2^ = 0.38). Thus, repeated stimulation made the middle-aged brains more vigorous by increasing their spike frequency in a consistent fashion across the whole length of the fictive motor program. In old brains, however, we were unsuccessful in fitting a singular non-linear curve to all the experimental graphs post-stim. We found that separate one-phase decay graphs best fit the curves for stim 1 (r^2^ = 0.29) and stim 2 (r^2^ = 0.26), while separate two-phase decay graphs best fit for stim 3 (r^2^ = 0.40), stim 4 (r^2^ = 0.52), and stim 5 (r^2^ = 0.55). Therefore, old brains are becoming less active in inconsistent ways across the entire period of the motor program, suggesting deeper deficits in network functionality with aging.

These results demonstrated that in middle-aged *Aplysia*, repeated stimulation produced a sensitization-driven increase in network activity across trials, which we expect to correlate to increased distance travelled during the motor program. With age, the effect is flipped – repeated stimulation produces a habituated decrease in network activity. This reduced response from the aged network is likely the result of CNS aging, and the reduced output from aged brains to a stimulus may explain the decline in galloping response observed in aged animals.

### Aging also reduces the activity and rhythmicity of resting efferent neurons

The reduced average neuronal activity in aged brains during motor programs made us consider other properties of the efferent neurons that are also declining with age. Aging is also known to reduce neuromodulator expression in *Aplysia* [37] and alter sensory and motor neuron excitability in a different *Aplysia* behavior [14, 17], which could result in a general reduction in pedal efferent neuron activity, before and after a stimulus. A recent study in middle-aged *Aplysia* that performed nerve recordings in intact animals also identified slow waves of activity at rest in pedal nerve 10, a critical pedal efferent nerve that is considered a readout of locomotion [32], suggesting the presence of a background rhythm in middle-aged resting pedal efferent neurons, likely a slow background precursor to the behavior. Does aging make these pedal neurons less active at rest and diminish or weaken the parameters of this resting rhythm? If so, that could contribute to the reduced fictive locomotion response we observed previously. To determine if aging reduced the resting activity of these pedal neurons, we prepared *Aplysia* pedal ganglia for VSD recording and left them undisturbed in the imaging chamber for 1 hour, before recording spontaneous activity for 5 minutes in middle-aged (n = 7) and old (n = 7) isolated brains, with no stimulus-elicited motor program.

We first compared average neuronal activity during these spontaneous recordings between middle-aged and old brains as in the previous section, by comparing the average number of spikes per cell during a VSD recording (**Fig. 4A-B)**. We found that the # of spikes/cell for middle-aged brain spontaneous activity was significantly greater than that of old brains (t-test, middle-aged: 121.3 ± 2.4 vs. old: 103.7 ± 2.6, *p* = 0.0003) (**Fig. 4C)**. This reduction of spontaneous activity at rest indicates a generalized effect of aging on the activity of the efferent neurons.

**Figure 4.**
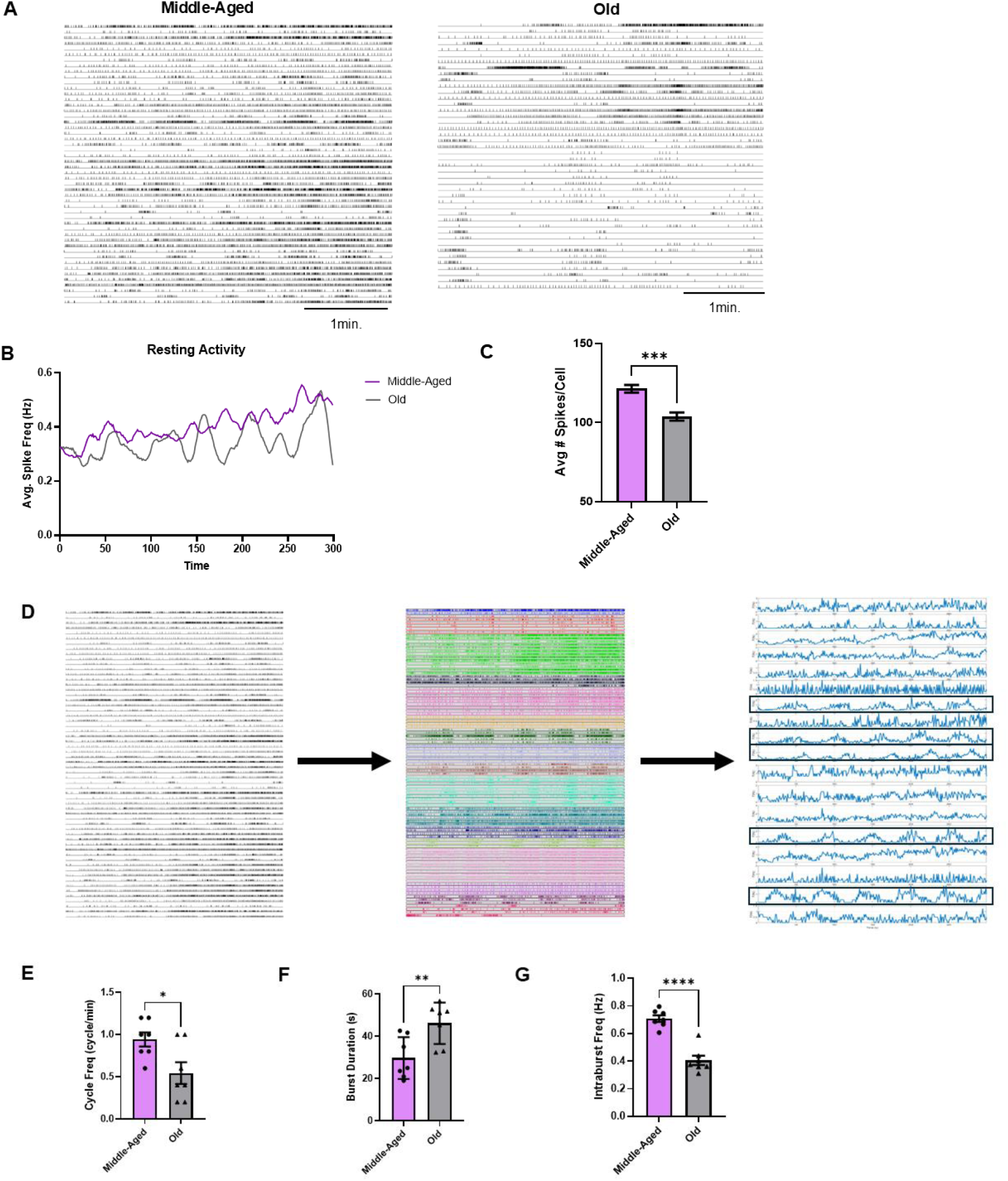
Aging reduces resting state activity in the pedal ganglion. **A)** Examples of spike rasters from VSD imaging in middle-aged and old pedal ganglia after 1 hour of rest. Middle-aged pedal neurons demonstrate noticeably more activity at rest compared to old pedal neurons. **B)** Average spike frequency graphs of all recordings of middle-aged (n = 7) and old (n = 7) isolated brains at rest. Rolling 10-s average plotted. **C)** Average spikes/cell of resting state activity shows a significant decrease from middle-aged to old (*** for *p* < 0.001). **D)** Workflow for characterizing resting rhythm. Spike trains of neurons (left) underwent unsupervised consensus clustering, producing statistically correlated functional clusters (middle). Average spike frequency is determined for each functional cluster (right) and power spectral density was used to identify which clusters are rhythmic (noted with black boxes) for calculation of parameters shown in **E-G. E)** Cycle frequency of resting rhythm decreases significantly in old brains (* for *p* < 0.05). **F)** Burst duration of resting rhythm bursts increases significantly in old brains (** for *p* < 0.01). **G)** Intraburst frequency decreases significantly in old brains (**** for *p* <0.0001).

**Figure 5.**
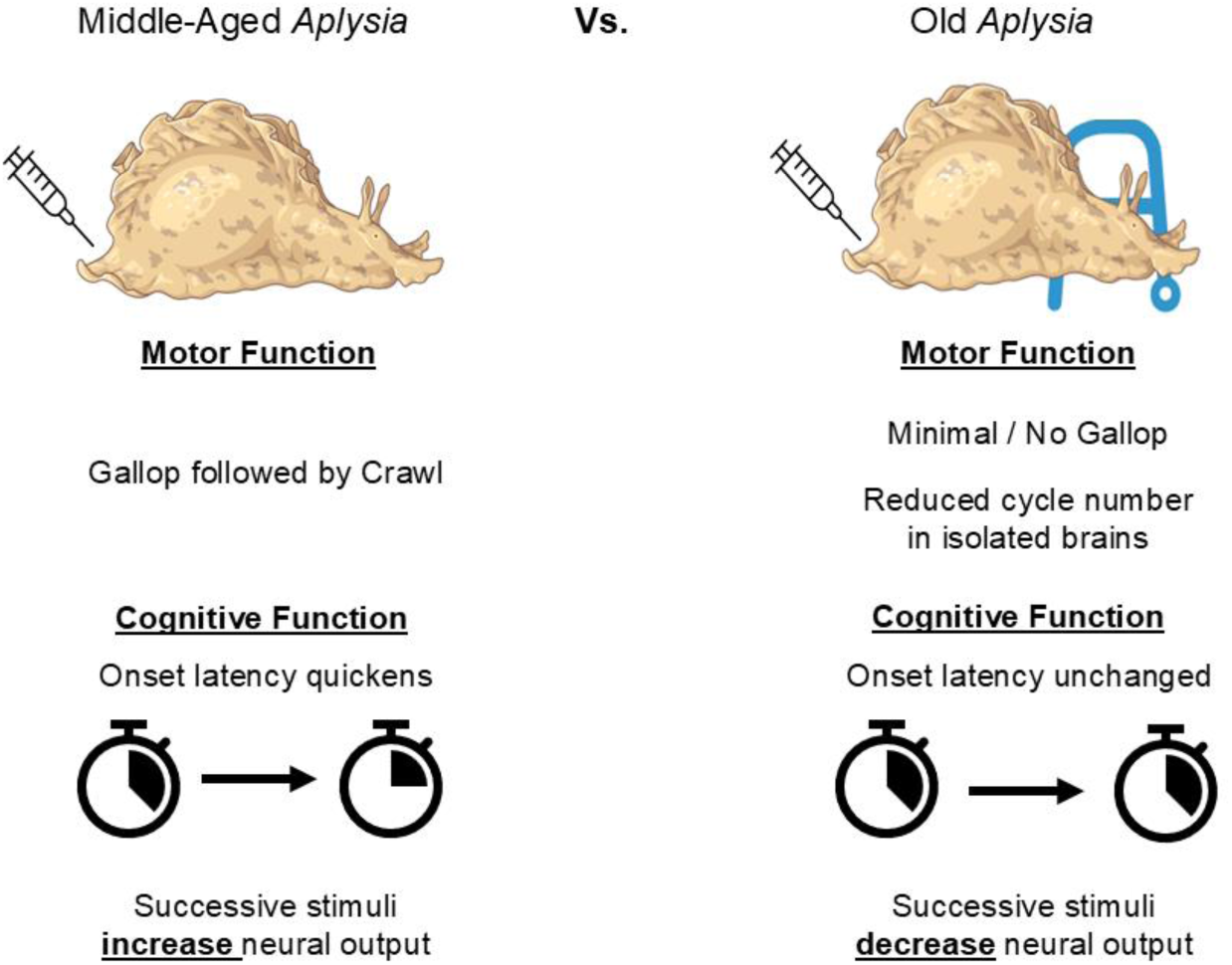
Summary of how aging affects *Aplysia* locomotion. Aging produces cognitive deficits in both intact animal behavior and isolated brain activity. Older *Aplysia* lose the ability to shorten initial head reach latency, and show a significant reduction in galloping. In their brains, we observed reduced neural activity at rest and after successive stimuli. We also saw reduced cycling capacity in isolated brains, suggesting aged animals may be more reliant on sensory-motor feedback from the periphery than middle-aged animals.

Our VSD imaging detected the presence of a slow bursting rhythm in the resting activity. To determine if this resting rhythm was diminished in aged preparations, we first developed a methodology for discriminating rhythmic and non-rhythmic neurons. We used a previously published unsupervised clustering algorithm [26] to pair up neurons into identified functional clusters and then used power spectral density to determine if these functional clusters were rhythmic (**Fig. 4D)**. If a cluster was rhythmic, the cycle frequency, burst duration, and intraburst frequency for the whole cluster was then calculated and averaged for each recording, allowing us to quantify this resting rhythm in middle-aged neurons and see if it was affected by aging.

Unpaired t-tests identified significant decreases in cycle frequency (middle-aged: 0.94 ± 0.08 cycles/min vs. old: 0.54 ± 0.13 cycles/min, *p* = 0.02), burst duration (middle-aged: 29.7 ± 3.7s vs. old: 46.2 ± 3.7s, *p* = 0.009) and intraburst frequency (middle-aged: 0.71 ± 0.02 Hz vs. old: 0.40 ± 0.03 Hz, *p* < 0.0001) in old brains compared to middle-aged brains (**Fig. 4E-G)**. Thus, this slow resting rhythm observed in middle-aged *Aplysia* pedal neurons is significantly weaker in older brains, representing another feature of aging in the neurons that participate in the motor network.

These results demonstrated a significant weakening of resting activity in old *Aplysia* pedal neurons, both in overall activity and with respect to a slow spontaneous bursting rhythm that may or may not correspond to slow crawling. This broader reduction in activity in pedal neurons at rest suggests that aging is broadly affecting the constituent members of the locomotion network. The reduced activity post-stimulus in aged brains is therefore likely produced by a combination of age-related decline in the CPG and the pedal neurons.

## DISCUSSION

As aging produces complex, heterogeneous decline in behavior and memory formation, exploring behavioral and cognitive aging in simple model systems presents opportunities to draw direct connections between age-related changes in neural activity to behavioral and cognitive loss. In this study, we show for the first time how aging affects motor function and memory formation within *Aplysia* escape locomotion. These findings lay the foundation for future studies to investigate how to mitigate and rescue age-related decline in this behavior, which may reveal insights about aging that are generally relevant across species.

### Behavioral and cognitive age-related decline emerge within specific components of *Aplysia*’s escape behavior

We began this study with an effort to characterize how non-associative learning in *Aplysia*’s escape behavior is changed by aging. However, the two main forms of non-associative learning, sensitization and habituation, can be encoded differently into the same network depending on stimulus interval and intensity [38]. Work in *C. elegans*’ reversal response has demonstrated varied responses to habituation training depending on inter-stimulus interval [39]. In the nudibranch mollusk *Tritonia diomedea*’s escape response, the same behavioral parameters can sensitize or habituate differently depending on stimulus intensity and interval [40]. In the same set of animals, one parameter of *Tritonia’s* escape behavior (onset latency) can even remain sensitized as another component of the escape behavior (cycle number) habituates [41]. In our experiments here, we studied three different parameters of the *Aplysia* escape behavior: head reach or onset latency, cycle number, and gallop share of all cycles. The only parameter of these in middle-aged *Aplysia* that underwent either sensitization or habituation was onset latency, which sensitized after an initial stimulus and remained sensitized with repeated stimuli. It is possible that a different stimulus protocol could produce different results, and further work is needed to identify other behavioral parameters of this behavior that may also be affected by learning. In our elderly animals, by comparison, we observed a loss of onset latency sensitization, the first identified site of cognitive aging within this behavior. While cycling remained robust in these aged animals, their galloping response was largely abolished. Thus, within this behavior, one parameter underwent cognitive aging, another underwent behavioral aging, and another is unaffected by aging.

Distinct impacts from aging on various components of the same network are consistent with a larger body of work demonstrating the heterogeneity of aging in behavioral circuits, even in simple invertebrates. For instance, while aging causes impaired graded response and faster habituation in *Aplysia*’s gill-siphon withdrawal reflex, it has no effect on gill respiratory pumping [24]. This difference was suggested to be the result of increased usage of the respiratory pumping pathway under normal conditions, or its necessary role in breathing. As crawling locomotion is a necessary behavior that *Aplysia* perform frequently and spontaneously [32], this may explain why crawling remains robust in aged animals.

These differences in behavioral aging may also be the result of different neurons within the network aging differently. In the freshwater mollusk *Lymnaea stagnalis*, aging causes feeding behavioral deficits through changes to specific synapses in the network [42]. Even within a single neuron of this network, aging has been shown to differentially affect synaptic connections [43]. In *Aplysia*, two different cholinergic neurons of the gill-siphon withdrawal reflex show distinct age-related changes to their genomes, including to genes responsible for neuronal function and synaptic plasticity [44]. If the neurons responsible for the various elements of the locomotion behavior are aging differently, this could explain why some parameters are affected by aging when others are age-invariant.

While aged *Aplysia* did not undergo onset latency sensitization, we found it interesting that their onset latency was already quick, compared to the middle-aged *Aplysia* which started off slower (**Fig. 1B**). In our isolated brain experiments, old animals also did not quicken their latency, but they started off slow and remained slow even with repeated stimulation (**Fig. 2B**). We speculate that in our behavioral experiments, our middle-aged *Aplysia* adapted to the new context of the testing arena and were in a state of rest during delivery of the first stimulus, while the old *Aplysia* were not. *Aplysia* are known to sleep [45], and sleep deprivation or disruption has been shown to interfere with learning in other *Aplysia* behaviors [46, 47]. Age-related sleep disruption has been demonstrated in many species [48] and has been demonstrated to contribute to cognitive loss in elderly humans [49]. One possibility is thus that the ability of our aged animals to rest is disrupted by the change in context when they are moved into the testing area, resulting in a quickened latency in intact animals. Deafferentation, however, eliminates this confound, and reveals the extent of CNS cognitive and behavioral aging via a slower latency in isolated brains that are unable to quicken with repeated stimulation.

### Differences in behavioral and cognitive aging between isolated brains and intact animals

One central advantage of utilizing marine invertebrates for studying the neural basis of behavior and learning has been that their behavior can be reliably reproduced in isolated or semi-intact preparations. These behaviors often utilize small neuronal populations and do not require the entire body to be generated with the correct pattern and period. The gastric/pyloric rhythm in crustaceans [50], the escape swim of *Tritonia* and *Pleurobranchaea* [51], and the escape response of medicinal leech [52] are all examples of simple behaviors in invertebrates that have been robustly interrogated in isolated preparations precisely because the fictive network activity in their neurons closely mirrors the real behavior of the intact animal. *Aplysia* has made numerous contributions to this field of work as well – *Aplysia*’s gill-siphon withdrawal reflex [53], tail withdrawal reflex [54], feeding motor program [55], and escape locomotion [28] have all been fictively generated in isolated preparations with time courses and activity patterns that are consistent with what is observed in the intact animal, supporting decades of work exploring the neural basis of these behaviors as well as different forms of learning within them.

We sought to determine whether the behavioral and cognitive aging we observed in our intact animal experiments persisted in isolated brains, as aging is also known to affect other body systems such as the muscles [1]. We observed cognitive aging in one parameter of isolated brains – onset latency of the locomotion motor program could sensitize in brains from middle-aged but not old animals. Notably, onset latency sensitization in middle-aged brains was more transient than in middle-aged animals, suggesting a peripheral contribution to persistent sensitization.

While we also observed no change in cycling with repeated stimulation within either age group, we did observe an overall reduction of cycling in all trials in aged brains compared to middle-aged brains (**Fig. 2C&E)**. This behavioral aging was specific to the CNS, as we did not observe this in intact animals.

This CNS aging may represent a network-level signature of age-related dysfunction that has yet to manifest in the intact behavior, likely due to the emergence of compensatory mechanisms. It is possible that the peripheral nervous system (PNS) is playing a role in maintaining locomotion cycling in aged animals. The role of the PNS in regulating behavior and learning has been previously explored in marine invertebrates. The PNS has been demonstrated to make critical contributions to the development of and persistence of habituation in *Aplysia*’s gill-siphon withdrawal reflex [56–59]. The nudibranch mollusk *Berghia stephanieiae* utilizes both CNS and PNS processes to produce a defensive posturing behavior known as bristling, and the PNS is able to override the CNS under certain circumstances [60]. The interactions between octopus brain and arms have produced a robust model of a distributed network where the CNS and PNS work synergistically to generate behavior, a framework for embodied intelligence [61–64]. It is possible that the PNS may play a role in both normal behavioral function and learning in *Aplysia* escape locomotion.

Aging has been shown in both invertebrates and vertebrates to affect the interactions between the CNS and PNS. In *Lymnaea,* semi-intact lip-CNS preparations of the feeding network have identified age-related decreases in CNS output, but not in PNS input – demonstrating that aging specifically affected central but not peripheral processing [65]. Aging has also been shown to affect the level of central versus peripheral control of habituation learning in *Aplysia*’s gill- siphon withdrawal reflex [66]. In both mammalian animal models and in human studies, there is growing interest in the role of the autonomic nervous system (ANS) in typical brain aging and utilizing ANS variability to predict neurodegenerative diseases [67, 68]. Future experiments utilizing semi-intact preparations of the *Aplysia* escape behavior may have the potential to elucidate the relative contributions of the PNS and CNS in producing regular behavior.

### Aging diminishes and disrupts network activity, before and after a stimulus

Simple marine invertebrates possess large neurons with easily accessible somata that make them very amenable to large-scale activity imaging with voltage-sensitive dyes [69]. Technical advances have enabled the ability to capture the voltage activity of dozens of neurons with single-cell, single-spike resolution [36]. VSD imaging has facilitated investigations into how neural networks produce behavior in multiple invertebrate species, including *Tritonia* [70], *Berghia* [35], crabs [71], and medicinal leeches [72]. VSD imaging has also been used to reveal novel network-level properties of learning within the *Tritonia* escape swim [73]. In *Aplysia,* VSD imaging has facilitated investigations into the networks that produce the gill-siphon withdrawal reflex [74], feeding [75], and escape locomotion [26, 27]. We sought to apply this technique to study how aging affects the locomotion network’s ability to reorganize itself during non-associative learning.

In middle-aged isolated brains we observed a steady increase in the intensity of the fictive motor program with each successive stimulus (**Fig. 3E**). Since we did not observe an increase in the cycle number in these recordings, this suggests that each cycle performed was more intense, which seems likely to translate to an increase in distance travelled over time. This would be consistent with a prior study of *Aplysia* escape locomotion sensitization, which found an increase in distance travelled after a sensitizing stimulus [33]. In our VSD recordings from old brains, by comparison, we observed a steady decrease in motor program activity with each successive stimulus. This decline in population activity represents a network component of cognitive aging, with sensitization giving way to habituation as animals age. We would also expect this to result in reduced distance travelled, but further work is needed to confirm this hypothesis.

How does repeated stimulation produce these increases in activity in youth, and why does it produce decreases in activity with old age? One potential explanation is serotonin modulation of both the CPG and the efferent neurons. Prior work has demonstrated that one of the primary modulators of the locomotion behavior is serotonin [76, 77], and many of the pedal efferent neurons possess 5-HT_1A_ and/or _2A_ receptors [78]. Delivery of serotonin precursor (5-HTP) is also sufficient to elicit spontaneous locomotion in middle-aged animals [79]. Sensitization learning in different *Aplysia* behaviors utilizes serotonin modulation to modify intrinsic properties, synaptic strength, neurotransmitter release, and tonic firing [80, 81]. However, dopamine is also a prominent neuromodulator of this motor program that is responsible for transitioning the network from gallop to crawl, and many pedal neurons also possess D_1-4_ receptors [25, 82]. We therefore speculate that repeated activation of this motor network is making multiple modulatory changes via serotonin and dopamine to the CPG and efferent neurons that ultimately result in a more vigorous escape motor program. This argument is supported by our demonstration of reduced vigor in the aged network with repeated stimulation. Aged *Aplysia* are known to have lower serotonin expression levels and reduced expression of dopamine receptors in pedal neurons [37, 82], so it is possible that this reduced response vigor in aged brains is the result of decreased or disrupted neuromodulation. It is also possible that aging causes a reduction in CPG neurotransmitter release onto the efferent neurons. Habituation in *Aplysia*’s gill-siphon withdrawal reflex been demonstrated to be produced in part by reduced transmitter release [83]. While the middle-aged escape response does not habituate, it is possible that habituation develops with age due to the reduction in modulator expression. It is also possible that if we extended the training beyond 5 trials, habituation may be observed.

However, we also observed differences in how network activity post-stimulus changed based on age. While middle-aged brains increased their output with each successive stimulus, they did so in a consistent fashion over time, such that a single two-phase decay curve was sufficient to fit every post-stimulus activity graph. We hypothesize that this is because the pedal efferent neurons have become more intrinsically excitable as a population, such that the same amount of CPG output produces a consistently stronger efferent response. Learning induced changes in excitability have been explored in other invertebrates – the mollusk *Hermissenda crassicornis* alters the excitability of visual-vestibular photoreceptors during associative learning [84, 85].

Altering the excitability of a network during learning has also been observed in various mammalian brain regions, and as such represents a conserved mechanism for learning-induced plasticity of a network [86]. Further work is needed to test this hypothesis, and to determine what cellular mechanisms may be potentially involved.

Aging has been shown to reduce intrinsic excitability and synaptic input to a critical motor neuron in the *Aplysia* gill-siphon withdrawal reflex [21, 87], and decrease intrinsic excitability and synaptic strength in mammalian circuits [88, 89]. As a result, we were not surprised to see that the aged network responded less intensely to an initial stimulus and became less intense with repeated stimulation. However, repeated stimulation reduced neural activity in an inconsistent fashion – a different decay curve was needed to fit each post-stimulus graph. This suggests that in addition to potential reductions in neurotransmitter tone and reduced excitability of aged efferent neurons, there are other disruptions to the aged network occurring with repeated stimulation that disrupted the activity post-stimulus. This may suggest that aging is also interfering with deeper components of the network we are not appreciating simply by looking at spiking activity. Future work is needed to identify how else the network’s response to repeated stimulation is disrupted in aging.

In addition to changes in neuronal activity, aging is known to affect many different biological and physiological functions [1, 2]. In *Aplysia*, aging has been shown to disrupt second messenger signaling and cellular metabolism in tail sensory neurons [19, 20, 90], likely contributing to behavioral and cognitive aging in its tail withdrawal reflex. We therefore utilized our VSD imaging technique to determine if aging altered the activity of pedal neurons at rest. This would allow us to indirectly assess the degree to which reduced fictive locomotor activity is the result of specific deficits in the CPG versus deficits across the efferent neurons as well. We did observe a decrease in network activity at rest in old pedal ganglia compared to middle-aged pedal ganglia (**Fig. 4A**). However, the more dramatic decrease we observed was a significant weakening of the slow oscillatory activity we observed in many middle-aged pedal neurons at rest. This slow oscillation, which we term the “resting rhythm”, has been observed in nerve recordings in middle-aged intact animals [32], and is present in many neurons that burst rhythmically during the motor program. While we do not currently know the precise function or source of this resting rhythm, its weakened state in aged brains suggests further deterioration within the CNS that may affect the fictive locomotion motor program.

What is the role of this resting rhythm in the pedal neurons? We hypothesize it represents a slow background readiness state for the locomotion behavior that is disrupted in aging. This disruption could be driven by a combination of reduced upstream input as well as degraded intrinsic properties of pedal neurons. In humans, fMRI studies have identified a series of brain regions active at rest that are collectively referred to as the “default mode network” (DMN) [91]. While the role of the DMN remains a topic of investigation, arguments have been made about its role in cognition [92]. In older humans experiencing typical brain aging, the DMN has been shown to be reduced in several critical regions involved in learning, including the hippocampus [93]. The DMN is also further dysregulated in older patients with neurodegenerative diseases like Alzheimer’s disease [94]. However, the indirect nature of human neuroimaging studies has limited the ability to directly study or manipulate the DMN. Future work attempting to rescue age-related behavioral and cognitive decline in this behavior should consider measuring changes in this resting rhythm to determine if the effects are localized to the fictive behavior or are generally improving the resting state of the aged brain.

### *Aplysia* escape locomotion as a novel paradigm for exploring age-related decline of rhythmic behavior

Behavioral and cognitive aging have been observed across various species and continue to be of strong clinical relevance to humans. To effectively study how aging develops within the brain and develop interventions to mitigate and rescue age-related decline, however, there is a benefit to utilizing systems with simple nervous systems to more directly draw comparisons from brain activity to behavior. *Aplysia* escape locomotion is well positioned as a paradigm for continued exploration into behavioral and cognitive aging, as it is a multi-phase rhythmic network with complex modulation that is highly amenable to network-level analyses of neural activity. We have demonstrated here that aging affects both the escape behavior and the neural network that produces it in heterogeneous ways. These findings are necessary to inform future work investigating how the network is declining in age and what types of interventions can successfully rescue said decline. *Aplysia*’s escape behavior is well positioned to yield general insights into the fundamental mechanisms of aging in rhythmic behavior and to help determine if and how we can rescue behavioral and cognitive aging.

## ACKNOWLEDGEMENTS

We would like to acknowledge Benedict Erhabor and Arnav Bajpai for assisting with certain behavioral experiments.

## AUTHOR CONTRIBUTIONS

Conceived and designed research: VKM, WNF, performed experiments: VKM, DM, analyzed data: VKM, DM, prepared figures: VKM, drafted manuscript: VKM, edited and revised manuscript: VKM, WNF, approved final version of manuscript: VKM, DM, WNF.

